# Gene regulatory network analysis identifies key genes and regulatory mechanisms involved in acute myocardial infarction using bulk and single cell RNA-seq data

**DOI:** 10.1101/2021.08.26.457775

**Authors:** Jiaxin Luo, Lin Wu, Dinghui Liu, Zhaojun Xiong, Linli Wang, Xiaoxian Qian, Xiaoqiang Sun

## Abstract

Cardiovascular and cerebrovascular diseases are leading causes of death worldwide, accounting for more than 40% of all deaths in China. Acute myocardial infarction (AMI) is a common cardiovascular disease and traditionally divided into ST-segment (STEMI) and non-ST-segment elevation myocardial infarction (NSTEMI), which are known with different prognoses and treatment strategies. However, key regulatory genes and pathways involved in AMI that may be used as potential biomarker for prognosis are unknown. In this study, we constructed weighted gene co-expression networks for differential expressed genes between STEMI and NSTEMI patients based on whole-blood RNA-seq transcriptomics. Network topological attributes (e.g., node degree, betweenness) were analyzed to identify key genes involved in different functional network modules. Furthermore, we used single-cell RNA-seq data to construct multilayer signaling network to infer regulatory mechanisms of the above key genes. PLAUR (receptor for urokinase plasminogen activator) was found to play a vital role in transducing inter-cellular signals from endothelial cells and fibroblast cells to intra-cellular pathways of myocardial cells, leading to gene expression involved in cellular response to hypoxia. Our study sheds lights on identifying molecular biomarkers for diagnosis and prognosis of AMI, and provides candidate key regulatory genes for further experimental validation.

## 1. Introduction

The morbidity and mortality of acute myocardial infarction (AMI) is still substantial worldwide despite the widespread access to reperfusion therapy [1]. In 2017, about 10.6 million cases of myocardial infarction were reported worldwide contributing to hospitalization and mortality [2].

AMI is divided into ST elevation myocardial infarction (STEMI) and non-ST segment elevation myocardial infarction (NSTEMI). The clinical manifestations of acute myocardial infarction are chest tightness, chest pain, and dyspnea. The pathophysiological process of acute myocardial infarction involves lipid deposition, vascular endothelial dysfunction, plaque formation, arterial stenosis, plaque rupture, and thrombosis [3]. Major risk factors for atherosclerosis included age, hypertension, hyperlipidemia, diabetes, obesity, and lack of exercise. The acute myocardial infarction is a multi-stage and multi-step disease, and patients with AMI are of significant heterogeneity, which make it difficult for clinicians to identify biomarkers and targets for prognosis and therapy [4].

Previous studies have demonstrated that discovering and controlling specific proteins can effectively control atherosclerosis. It is well known that lipid deposition is a key factor in atherosclerosis. Many experimental studies have shown that PCSK9 was associated with the phenotype of familial hypercholesterolemia. Based on this finding, the PCSK9 inhibitors (alirocumab and evolocumab) could dramatically decrease plasma LDL-C levels, even in patients who are taking the maximum dose of statins. PCSK9 inhibitors bring long-term benefits to patients with coronary heart disease, especially patients with myocardial infarction [5]. These approaches allow for the identification of novel gene, specific antibody and efficient therapeutic strategies. Inflammation is another key pathophysiological mechanism in atherosclerosis. In the CANTOS clinical trial (Canakinumab Anti-Inflammatory Thrombosis Outcomes Study), Canakinumab, an anti-IL-β monoclonal antibody significantly reduced the incidence of major adverse cardiovascular events (MACE) in the subjects with prior myocardial infarction [6].

The vascular endothelium is a thin layer of cells acted as a barrier to prevent lipid infiltration. Endothelial dysfunction is an independent risk factor of atherosclerosis. Numerous molecular and signal pathways are involved in the process of endothelial dysfunction, such as Nuclear factor kappa B(NF-κB) [7], endothelial nitric oxide synthase(eNOS) [8], AMPK-mTOR (AMP-activated protein kinase-mechanistic target of rapamycin kinase) signaling pathway [9]. However, the absence of reliable markers has hampered our understanding of specific role of endothelial dysfunction in atherosclerosis, especially myocardial infarction.

The cardiomyocytes (CM) were incapable of regeneration following injury in adult mammalian heart, but it can be regenerated for newborn mammalian heart. Several studies used RNA-sequencing to explore key molecular mechanisms of cardiomyocytes in myocardial infarction. Cui M et al. have recently described two factors, nuclear transcription factor Y subunit alpha (NFYa) and nuclear factor erythroid 2-like 1 (NFE2L1) transcription factors, that play a unique role in protecting against ischemic injury [10]. Ruiz-Villalba A et al. identified a unique subtype of cardiac fibroblasts (CF) that plays an essential role in ventricular remodeling process in response to cardiac damage [11]. The activated cardiac fibroblasts highly express collagen triple helix repeat containing 1 (Cthrc1) in the scar. Furthermore, it has been confirmed that the CTHRCI was a key regulator to heal scar process.

However, how different types of cells (e.g., endothelial cell (EC), CM and CF) interact with each other during the course of myocardial infarction are unclear. Furthermore, the key molecules involved in inter and intra-cellular signaling pathways between endothelial cell (EC), CM and CF remains poorly understood.

In this paper, to identify key genes and regulatory pathways involved in different subtypes of AMI, we collected both bulk and single-cell RNA-seq (scRNA-seq) data to construct weighted gene co-expression networks and multilayer inter-/intra-cellular signaling networks. Network module analysis, topological attribute analysis, functional enrichment analysis and scRNA-seq data analysis were performed. The results revealed that PLAUR (receptor for urokinase plasminogen activator) plays a vital role in inter- and intra-cellular signaling transduction of myocardial cells. Our study provided potential biomarkers for diagnosis and prognosis of AMI.

## 2. Materials and methods

### 2.1. RNA-seq data analysis

RNA-seq data was collected from Gene Expression Omnibus (GEO) database with accession number GSE103182 [12]. The RNA-seq data contained 30 samples of AMI patients, of which 15 were STEMI and 15 were NSTEMI. The raw data were corrected for unnecessary confounding variables and analyzed for differential gene expression with R package RUVSeq [13] and edgeR [14], respectively [12].

Based on the FPKM standardized expression matrix, we constructed the co-expression network for differential expressed genes using WGCNA R package [15] (version: 1.70-3). The main steps to construct co-expression network were as follows: (1) cluster the samples, and remove outlines; (2) select the soft threshold β; (3) construct co-expression network by ‘blockwiseModules’ function, which is used to divide genes into modules. The parameter ‘minModuleSize’ was set to 30. Other parameters not mentioned were default parameters in the WGCNA package.

For network visualization, we exported the co-expression network of gene modules (expect grep module), and then imported them into Cytoscape [16] software (version: 3.8.2), respectively. Also, we performed network topology properties analysis on every gene of every network by the ‘Analyze Network’ function in the Cytoscape software.

### 2.2. scRNA-seq data analysis

scRNA-seq dataset of heart tissue in human heart disease (coronary atherosclerotic heart disease and dilated cardiomyopathy) patients was derived from GEO database with accession number GSE121893 [17]. There were 25742 genes and 4933 cells in the data. Due to many missing values or dropouts in single-cell data usually, we made single-cell data imputation by scImpute [18] package before analysis. All parameters are default in the scImpute package.

We imported the scRNA-seq data into R4.0.3, and performed data analysis by Seurat[19] package (version: 4.0.0). First, ‘CreateSeuratObject’ function was used to create Seurat object, and ‘LogNormalize’ method in ‘NormalizeData’ function was used for data normalization. Then, we found 2000 highly variable genes through ‘FindVariableFeatures’ function. Next, we carried on dimensionality reduction by PCA method, based on the 2000 highly variable genes, which was scaled by ‘ScaleData’ function. The top 15 PCs was applied in analysis of cell clustering, by ‘FindNeighbors’ and ‘FindClusters’ function, whose resolution was set to 0.5. In addition, cell type classification was performed according to the marker genes described in the original literature[17]. Finally, UMAP method was used for visualization based on the first 15 PCs. Other parameters not mentioned were default parameters in the Seurat package.

### 2.3. Multilayer network construction

Based on the scRNA-seq data mentioned above, multilayer networks were constructed by our previously developed tool scMLnet [20] package (version: 0.2.0), which uses prior information and Fisher’s exact test to construct intercellular communication and intracellular transcriptional regulation network. We constructed multilayer networks with CM as receptor cells, macrophages (MP), EC, fibroblasts (FB) and smooth muscle cells (SMC) as ligand cells. All parameters are default in the scMLnet package.

### 2.4. Functional enrichment analysis

For further study, clusterProfiler [21] package (version: 3.18.1) was used for functional enrichment analysis in both RNA-seq and scRNA-seq data. In the WGCNA analysis of RNA-seq, in order to study the functions involved in each gene module, we performed gene ontology (GO) enrichment analysis on all genes of every gene modules respectively. In the multilayer network analysis of scRNA-seq, we extracted sub-network of specific gene from multilayer network. And GO enrichment analysis was employed in the downstream genes (transcription factors and target genes) of the sub-network. All parameters were the default parameters in the clusterProfiler package.

## 3. Results

### 3.1. Gene co-expression networks

Base on differential expression analysis, 323 differentially expressed genes (DEGs) were obtained, of which 180 genes were highly expressed in STEMI and 143 genes were highly expressed in NSTEMI (Figure 1) [12]. When clustering the samples, we found the sample 13003 was a outlier, thus we removed it from data for further analysis (Figure 2A-B).

**Figure 1.**
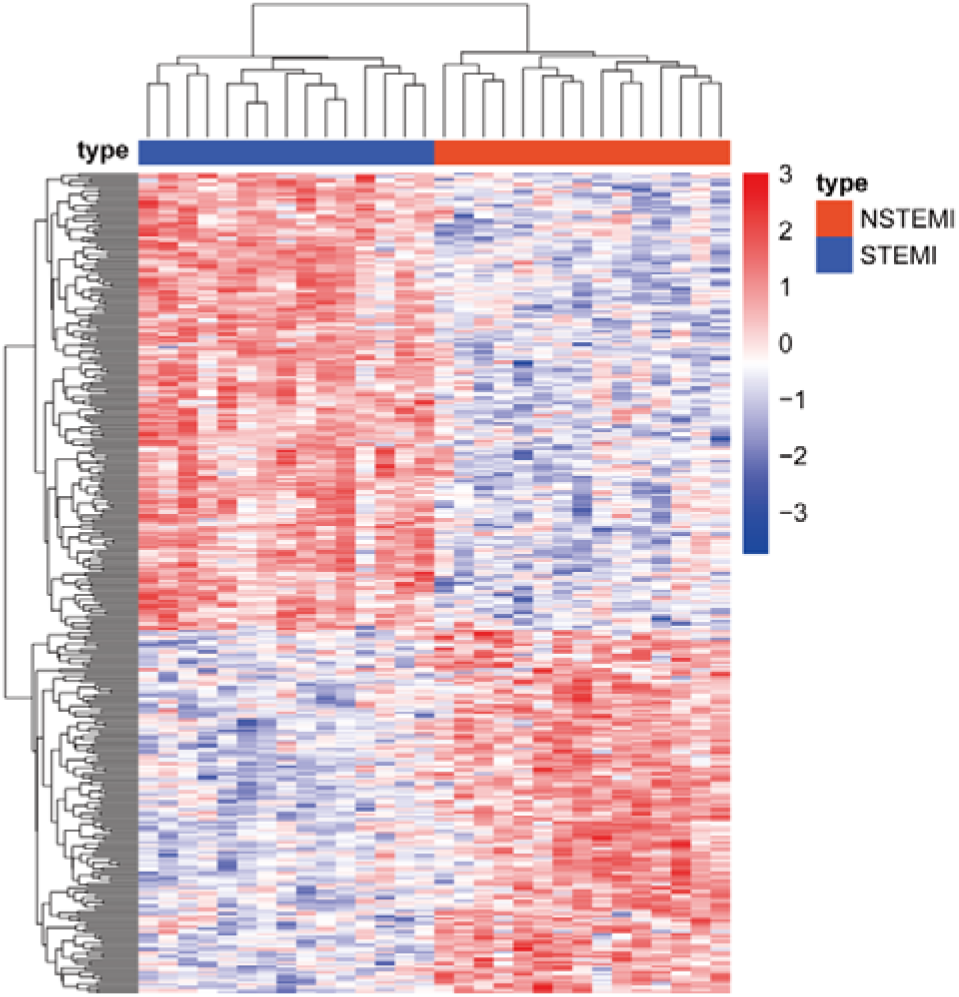
Heatmap of 323 DEGs between STEMI and NSTEMI.

**Figure 2.**
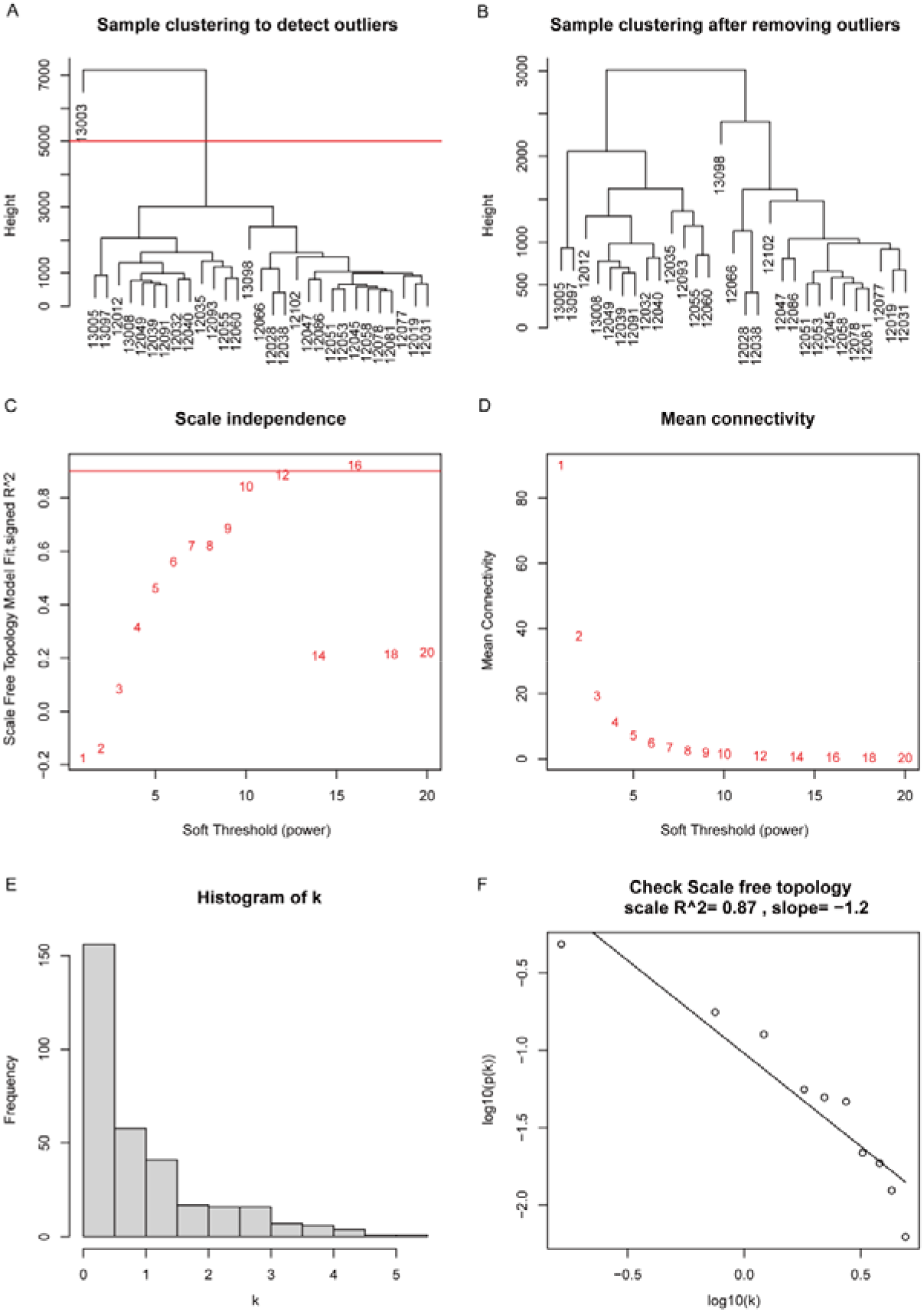
Co-expression network construction. (A-B) Clustered sample, and removed outlier samples. (C-D) Selected soft threshold β. (E-F) Check scale-free network.

The key step to construct co-expression networks by WGCNA is to select soft threshold β which take the adjacency matrix between genes β pow. The soft threshold β is used to make the co-expression network conform to the distribution of scale-free network. Because biological network has the characteristics of scale-free network, which means that the degree of nodes in the network obeys power-law distribution. In other words, it meets the negative correlation between log (k) and log [P (k)], where k represents the degree of nodes and P (k) represents the frequency of nodes with degree k. The larger the correlation coefficient R^2 between log(k) and log [P (k)], the more significant the characteristics of the scale-free network.

When β was 12, the correlation coefficient R^2 was greater than 0.9 (Figure 2C-D), and the network approached the distribution of scale-free network (Figure 2E-F). For reducing the operation time, we chose 12 as β, and then used ‘blockwiseModules’ function to construct co-expression networks. Finally, 323 DEGs were divided into 4 modules, each corresponding to each color (Figure 3A). Genes that didn’t belong to any of the four modules were classified into gray module.

**Figure 3.**
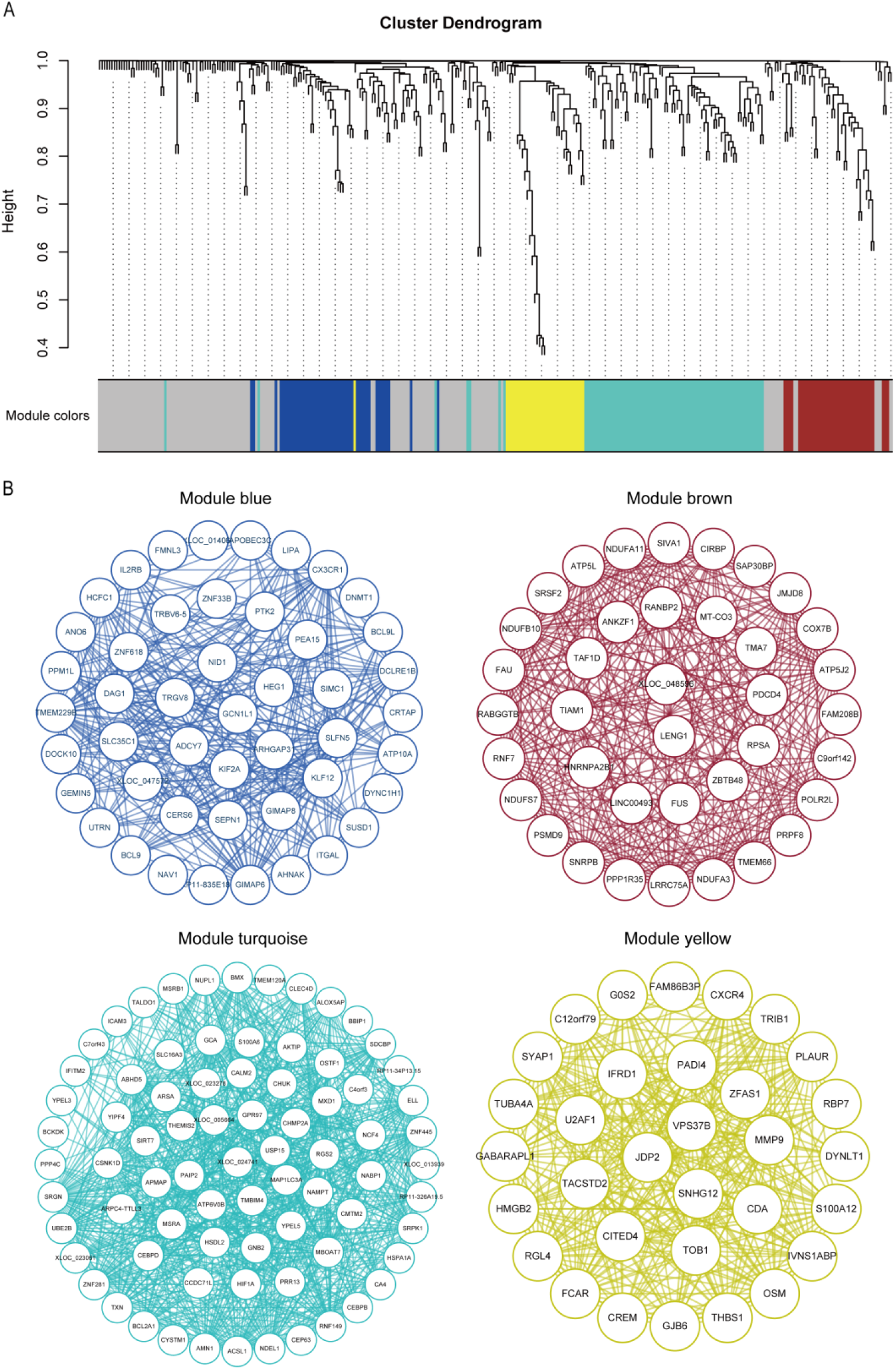
Co-expression networks visualization. (A) Cluster dendrogram and module division of genes. (B) Gene modules networks visualization by Cytoscape.

### 3.2. Network visualization and analysis

We imported the co-expression network of each module into Cytoscape software for visualization (Figure 3B), and used the “Analyze Network” function to analyze the network topology attributes of all nodes of each network. Degree is a count of the number of edges directly connected to a node in the network, and betweenness is defined as the proportion of the number of paths passing through the node in the total number of shortest paths in the network. That the greater the degree and betweenness of a node, means that the node plays an important role in the network and might be of great biological significance. Base on them, we identified potentially important genes in the modules (Figure 4). In the blue module, TMEM229B and GIMAP6 were two genes with high degree and betweenness; In the brown module, CIRBP had the highest betweenness, but its degree was slightly lower. While the degree of ATP5L, NDUFB10 and ATP5J2 were higher, but the betweenness of them was relatively low; In the turquoise module, RNF149, SRGN and ABHD5 had relatively high degree and betweenness; In the yellow module, although DYNLT1 had the highest betweenness, its degree was relatively backward. In addition, PLAUR, RGL4, IVNS1ABP and FCAR had relatively high degree and betweenness.

**Figure 4.**
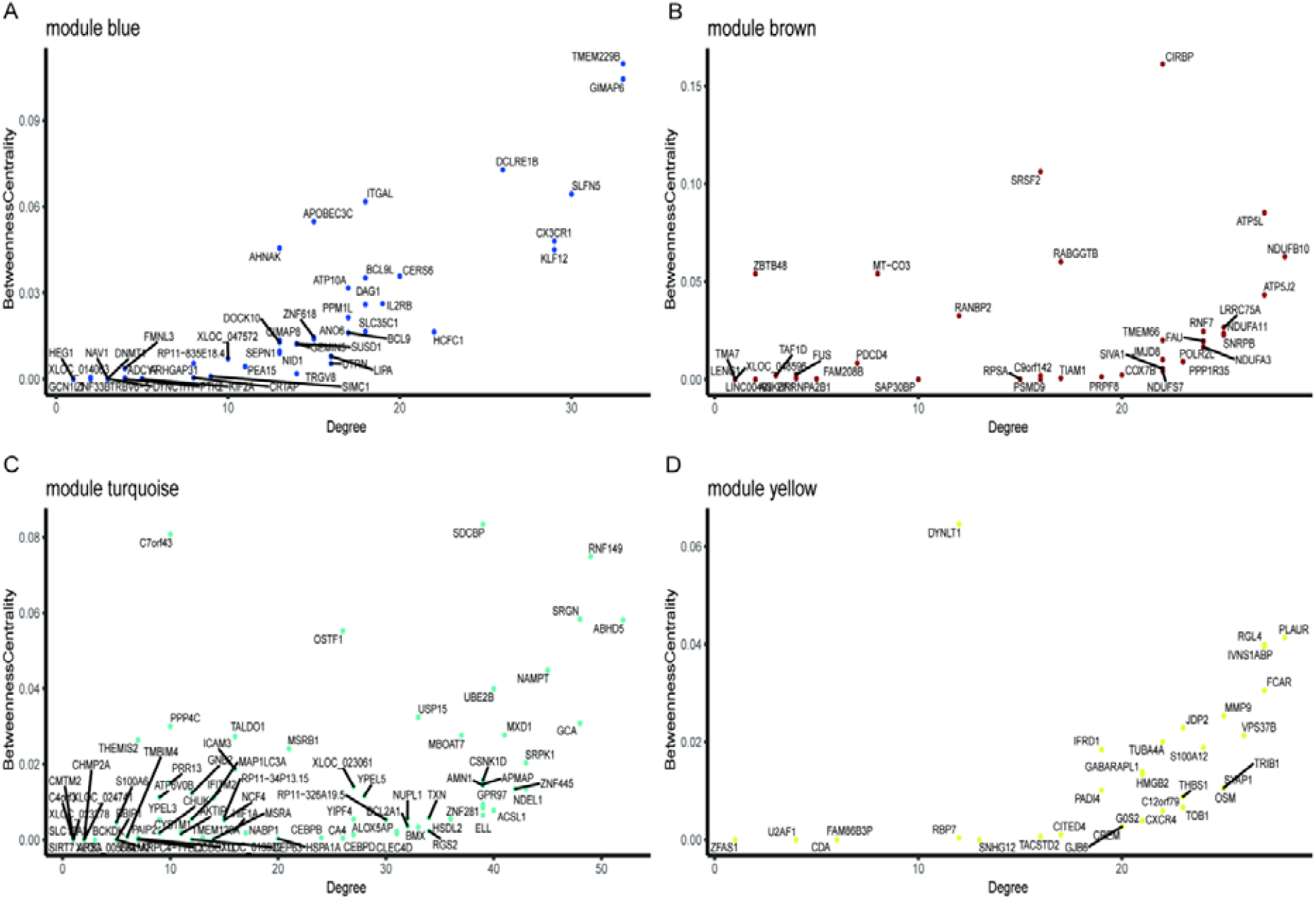
Scatter diagram of degree-betweenness centrality.

For further study of the biological processes involved in each gene module, we used the ‘enrichGO’ function in the clusterProfiler package to perform GO enrichment analysis on all genes of each gene module (Figure 5). We used “FDR” as the correction method of multiple hypothesis test, 0.05 as the threshold of P value and 0.2 as the threshold of Q value for enrichment analysis. The blue module was mainly significantly related to biological processes such as cell matrix adhesion and regulation of cell shape. Biological processes such as ATP metabolic process, RNA slicing and mRNA slicing were significantly enriched in the brown module. The yellow module was mainly significantly related to biological processes such as regulation of apoptotic signaling pathway, neural degradation and neural activation involved in immune response. However, no biological processes were significantly enriched in the turquoise module.

**Figure 5.**
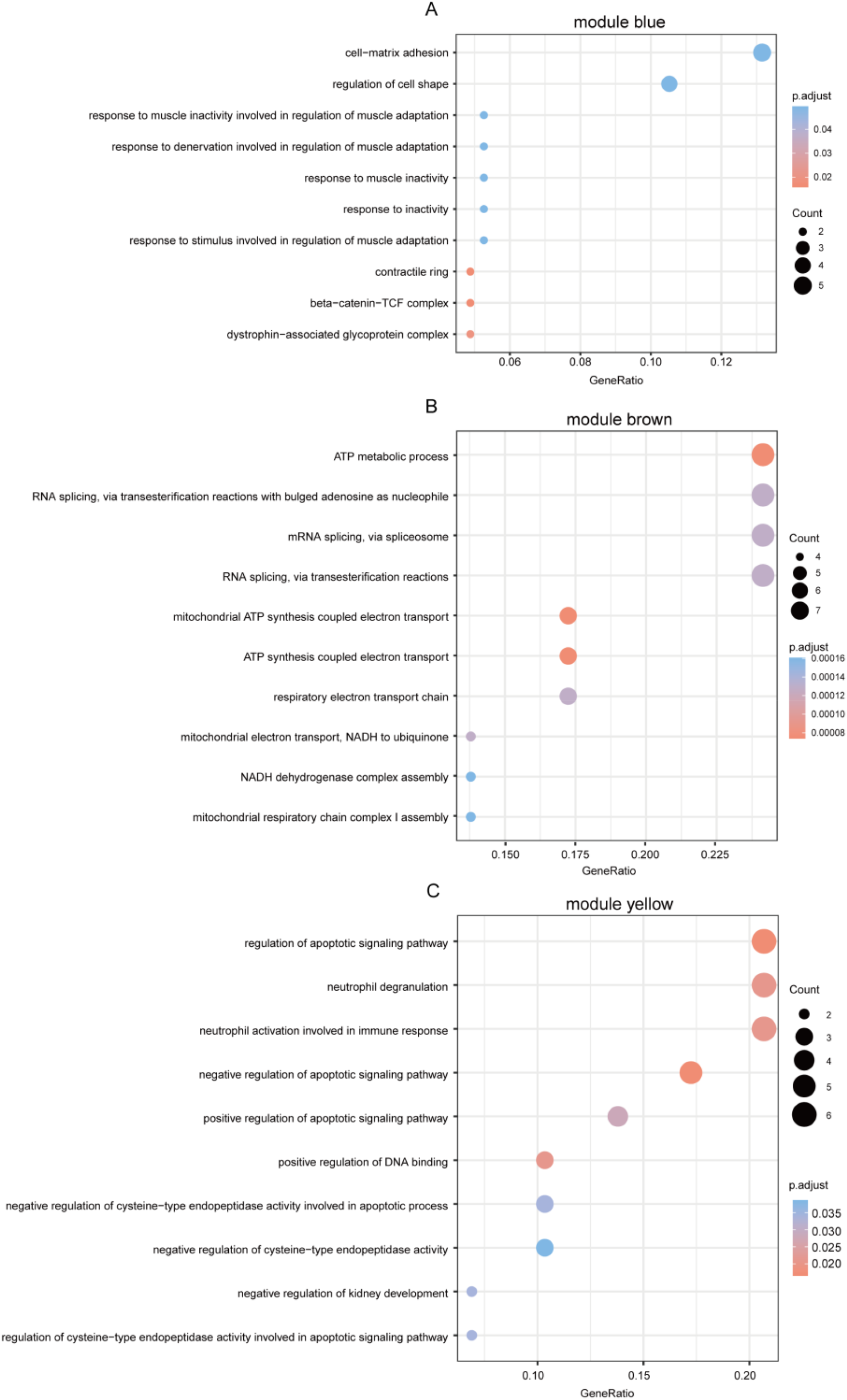
GO enrichment analysis for each gene module.

### 3.3. Multilayer networks

After data analysis of scRNA-seq, five cell types were identified, namely CM, MP, EC, FB and SMC (Figure 6). In addition, two clusters were not identified and were labeled as “UN1” and “UN2” respectively.

**Figure 6.**
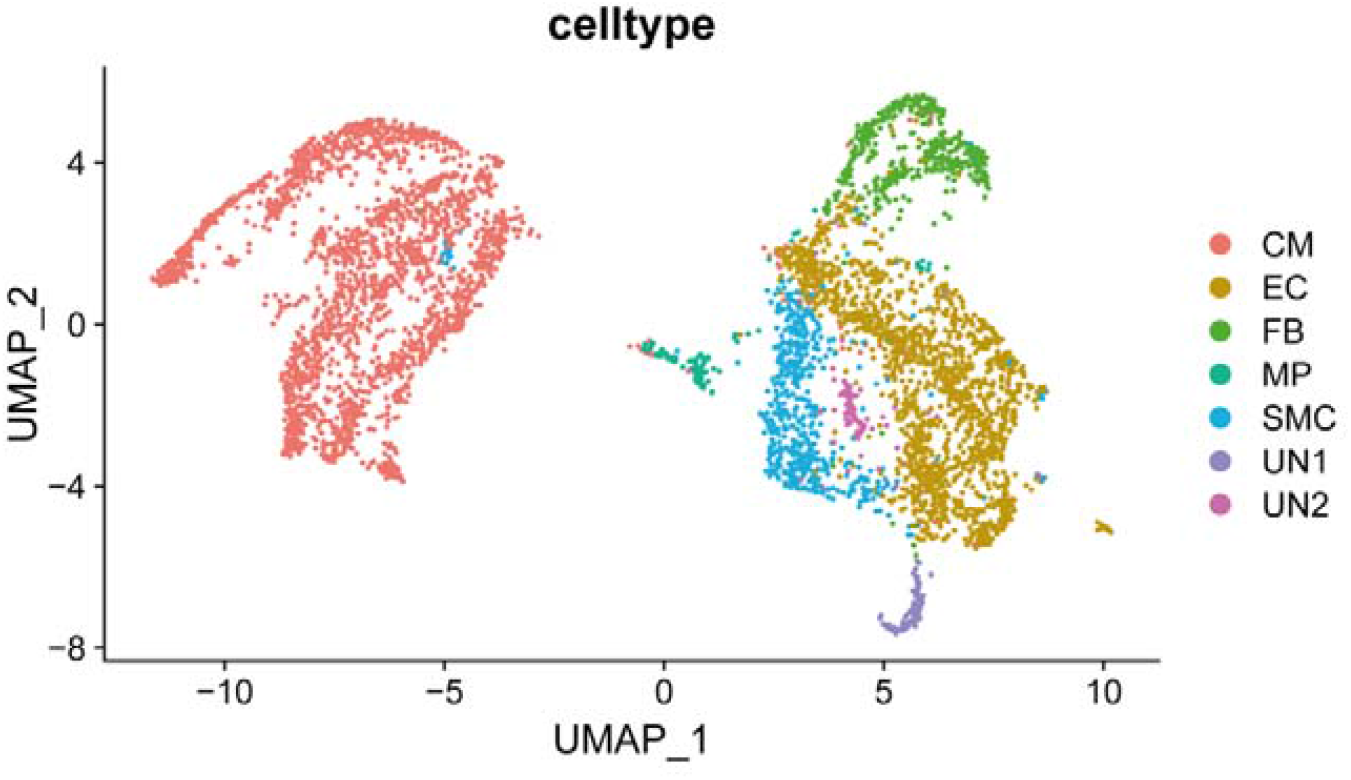
Cell-type clustering and identification

Then, we used CM as receptor cells and MP, EC, FB and SMC as ligand cells to construct multilayer networks respectively (Table 1, Figure 7). We found that PLAUR as a receptor existed in the multilayer networks of EC-CM and FB-CM (Figure 8A-B). Moreover, the downstream genes regulated by PLAUR receptor were the same in both networks. Therefore, we extracted the downstream genes regulated by PLAUR receptor for GO enrichment analysis (Figure 8C). We observed that downstream genes were significantly enriched in biological processes such as regulation endothelial cell proliferation, pri-miRNA transcription and cellular response to oxygen levels.

**Table1.** Result of multilayer networks.

| Receiver cell types |  | MP | EC | FB | SMC |
| --- | --- | --- | --- | --- | --- |
| Sender cell types |  | CM | CM | CM | CM |
| Number of pairs | Ligand-Receptor pairs | 13 | 150 | 188 | 63 |
|  | Receptor-TF pairs | 352 | 2183 | 1891 | 1184 |
|  | TF-TG pairs | 293 | 320 | 320 | 308 |
| Number of molecules | Ligands | 9 | 55 | 65 | 28 |
|  | Receptors | 13 | 84 | 73 | 43 |
|  | TFs | 85 | 99 | 99 | 95 |
|  | TGs | 87 | 87 | 87 | 87 |

**Figure 7.**
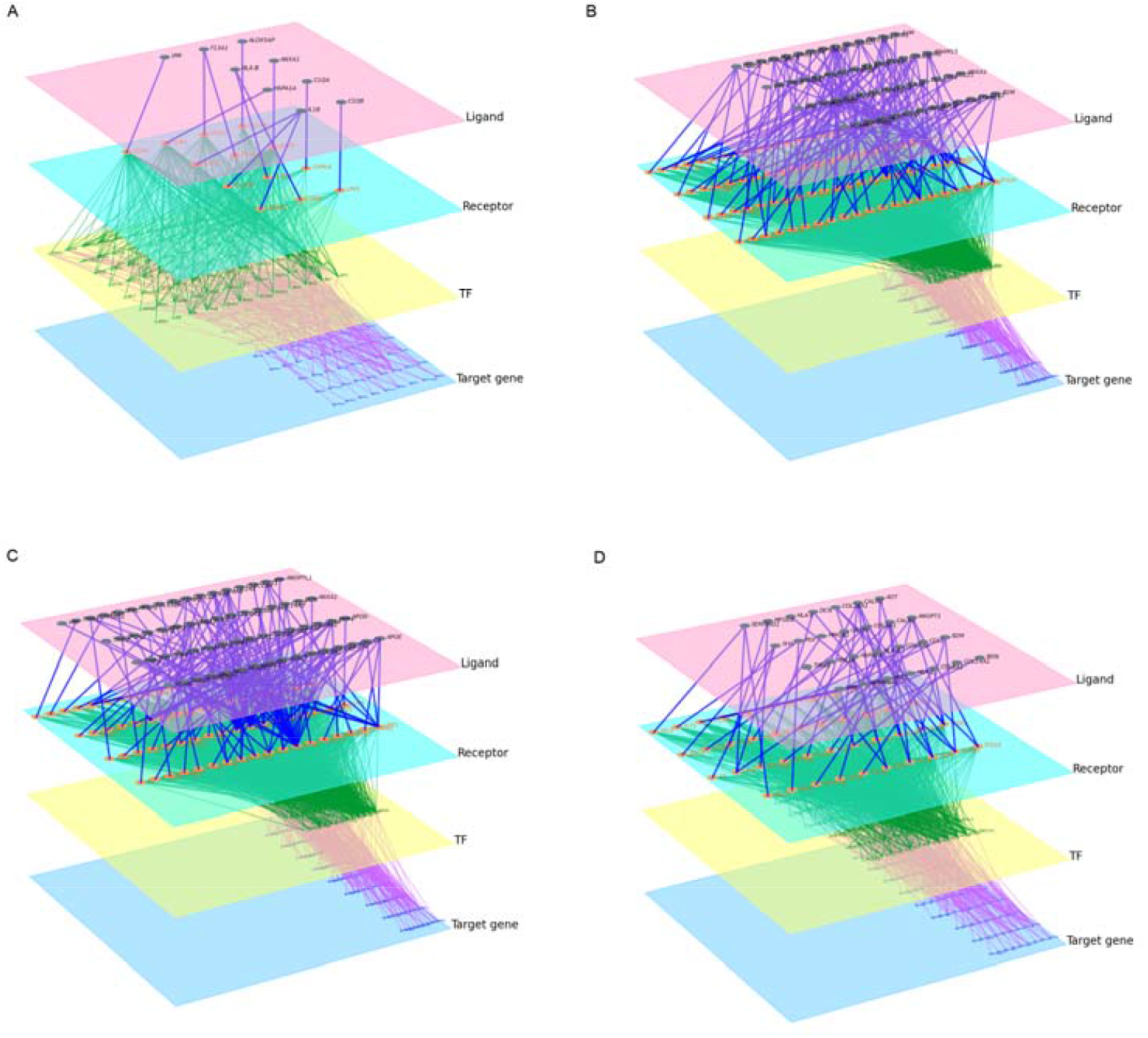
Multilayer networks visualization. Multilayer networks of MP-CM (A), EC-CM (B), FB-CM (C) and SMC-CM (D).

**Figure 8.**
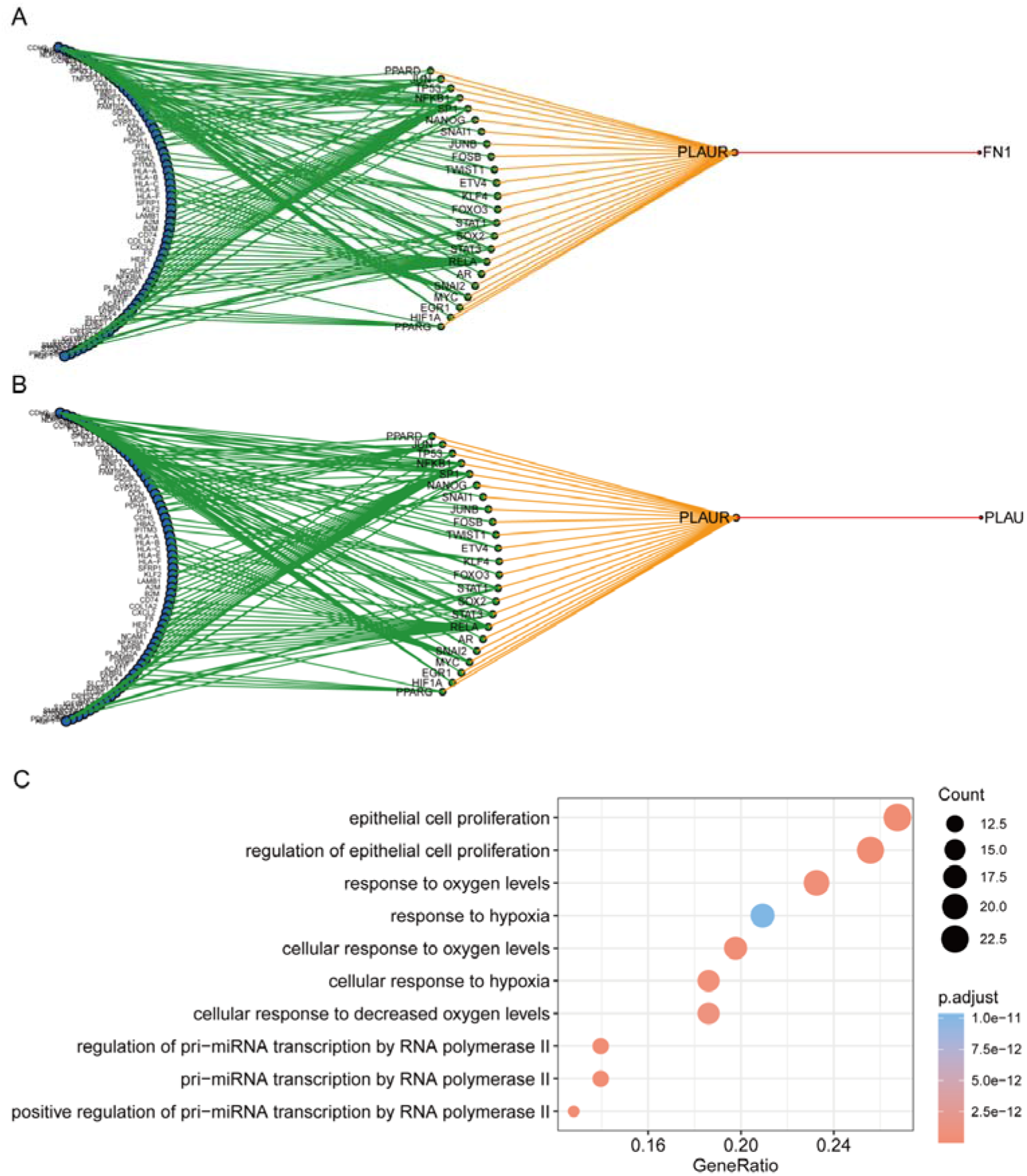
Visualization and analysis of PLAUR-related subnetworks. (A) PLAUR related subnetwork in the FB-CM multilayer networks. (B) PLAUR-related subnetwork in the EC-CM multilayer networks. (C) GO enrichment analysis for downstream genes regulated by PLAUR.

## 4. Discussion

Based on whole-blood RNA-seq transcriptomics, this study represents the weighted co-expression networks of differentially expressed genes between STEMI and NSTEMI patients. In addition to identify key genes involved in different network modules using network topological attributes, we revealed regulatory mechanisms of key genes by multilayer signaling network based on single-cell RNA-seq data. Furthermore, we identified PLAUR as a crucial receptor gene in transducing inter-cellular signals from endothelial cells and fibroblast cells to intra-cellular pathways of myocardial cells, leading to downstream gene expression involved in cellular response to hypoxia.

We mainly analyzed four modules of the co-expression networks constructed by WGCNA. According to GO enrichment analysis, those four modules were associated to some important biological processes, such as cell matrix adhesion, cell shape, apoptotic signaling pathway, neural degradation, neural activation, ATP metabolic process, RNA slicing and mRNA slicing. In the blue module, TMEM229B and GIMAP6 express highly. CIRBP, ATP5L, NDUFB10 and ATP5J2 had high betweenness centrality. RNF149, SRGN and ABHD5 were ranked as top genes in the turquoise module according to degree and betweenness centrality. In addition, PLAUR, RGL4, IVNS1ABP and FCAR were genes with relatively high degree and betweenness centrality in the yellow module, where although DYNLT1 had the highest betweenness centrality, its connectivity was relatively low. Previous study has proven that the process atherosclerosis involved in lipid metabolism, inflammatory, endothelial dysfunction, monocyte transendothelial migration and apoptosis. It has also been verified that TMEM229B is a novel gene which was highly expressed untreated islets and strongly suppressed by STZ. It means TMEN229b play a protective role in β-cell function [22].

In the multilayer networks of EC-CM and FB-CM, PLAUR was found as a crucial receptor that mediates inter-cellular and intra-cellular signaling pathways and regulates numerous downstream genes which are involved in endothelial cell proliferation, pri-miRNA transcription and cell response to oxygen levels. PLAUR is a glycosyl-phosphatidylinositol (GPI)-anchored membrane protein which is associated with cell signaling because it forms a multi-protein complex with neighboring tansmembrance receptors, such as EGFR [23]. Inhibition of PLAUR can inhibit tumor growth, invasion. PLAUR can be released into blood and be detected as a biomarker in small cell cancer [24], breast cancer [25]. In this study, we found that PLAUR had high degree and betweenness centrality, and involved in apoptosis signal pathway and immune response.

Nowadays in clinical practical, the major biomarkers of myocardial infarction are creatine kinase-MB (CK-MB) and cardiac troponin (cTn). The measurement of CK-MB and cTn is superior because of the high sensitivity and specificity for myocardial damage [26]. However, the limitation of CK-MB and cTn is obviously. CK-MB and cTn will elevate in skeletal muscle injury, marathon runners, chronic renal failure and hypothyroidism [27]. We expect that the key genes identified in this study represent promising novel biomarkers for myocardial infarction. Such biomarkers may facilitate timely diagnosis of AMI and identification of successful reperfusion after thrombolysis. Moreover, more and more drugs targeting a single gene or protein will have a great impact on decreasing of MACE. Amon them are PCSK9 inhibitor and anti-IL-β monoclonal antibody. The above-mentioned genes may be explored as potential targets for myocardial infarction.

In summary, in this study, we employed bulk and single-cell RNA-seq data to construct gene regulatory networks and cell-cell communication networks to identify key genes involved in AMI. Our study sheds lights on identifying molecular biomarkers for diagnosis and prognosis of AMI, and provides candidate key regulatory genes for further experimental validation.

## Data availability

The gene expression datasets as well as clinical information of the patients were downloaded from the NCBI GEO database, as described in the main text.

## Acknowledgements

X Sun was supported by grants from the National Natural Science Foundation of China (11871070), the Guangdong Nature Science Foundation (2020B1515020047).

## Author contributions

X.S. and X.Q. designed research; J.L performed data analysis; J.L. and L.W. analyzed data. J.L., L.W., D.L., and Z.X. involved in discussion. J.L., L.W., and X.S. wrote the manuscript. All authors proofread the paper.

